# Twelve species of human parasites make up half of the literature on microbial eukaryotes

**DOI:** 10.1101/2025.08.20.671258

**Authors:** Joanna A. Lepper, H.B. Beryl Rappaport, Angela M. Oliverio

**Author notes:** Corresponding author, Syracuse University Department of Biology, Life Sciences Complex, 107 College Pl, Syracuse, NY 13210.

## Abstract

Although microbial eukaryotes comprise the majority of eukaryotic phylogenetic diversity and inhabit nearly all ecosystems globally, most research focuses on only a few species of human parasites. Here, we quantify the extent of research on known microbial eukaryotic species. Nearly half of the mentions of protist species on PubMed referenced only 10 species included in the Protist Ribosomal Reference (PR2) Database. Likewise, although most samples in the PR2 database are free-living protists from aquatic environments, 12 species of human parasites comprise 47% of the literature. Research efforts that focus on only a handful of eukaryotic lineages severely limit our understanding of the fundamental biology of eukaryotic cells. We highlight recent efforts to characterize novel eukaryotic lineages that have resulted in a new understanding of the ‘rules of life’ and identify key lineages that are notably absent or limited in the literature, despite their abundance and significance across global ecosystems.

---

Although protists (e.g. microbial eukaryotes) comprise the vast majority of eukaryotic life on Earth^1^, with important contributions to global environmental systems^2,3^ and human health outcomes^4^, only a handful of lineages are the focus of most research efforts. Consequently, our understanding of fundamental ‘rules’ of eukaryotic life remain limited at best and misguided or incorrect at worst. Characterizing non-model eukaryotes, many of which are from poorly sampled clades or habitats, has repeatedly broadened our view on what is possible in biology^5,6^. For example, it was previously thought that nitrogen fixation was exclusive to prokaryotic cells, but recent research on a species of algae, *Braarudosphaera bigelowii*, led to the discovery of a new symbiotic organelle in eukaryotic cells, the nitroplast, that aids in nitrogen fixation^7^. Likewise, the discovery of non-canonical genetic codes in ciliates^8,9^ and other lineages of microbial eukaryotes^10,11^ challenges our understanding of what universal codons are assigned to code for specific amino acids. Although it is generally appreciated that only a few ‘model’ eukaryotic lineages are well-studied, we have lacked a quantitative understanding of the extent of this research bias.

In an effort to shine light on the bias within protist research, as many new clades and lineages are being recovered from ‘omics approaches^12,13^, we performed a quantitative assessment of the number of published articles across microbial eukaryotic species. Inspired by a recent parallel effort in bacteria^14^ which found that ten species out of over 43,000 made up half of literature, and 74% of bacterial species were left unstudied, we sought to quantify the extent of research bias in microbial eukaryotes. A quantitative understanding of key research gaps and current sampling and phylogenetic biases is a critical step in guiding future research efforts in microbial biology.

## Top protist species in literature

To quantify research efforts across microbial eukaryotes, we performed a bibliometric analysis of PubMed (*via* the PubMedR package). To determine the number of papers for each species of protist in the database, we searched for species names (genus and species) in the title or abstract. We used the <u>P</u>rotist <u>R</u>ibosomal <u>R</u>eference <u>D</u>atabase (PR2) to obtain a list of named protist species that also had corresponding 18S rRNA gene sequence data. This resulted in 8,456 unique microbial eukaryotic species which were mentioned a total of 242,844 times in PubMed as of April 11, 2025. We emphasize that our analyses here are limited to species with named taxonomies. The actual number of protist species is much greater, with estimates ranging from two to ten million or more species^15^, further underscoring the extreme lack of characterization across the eukaryotic tree of life. Although more protist species are denoted in NCBI (~60,000 species^16^) many lack species names and/or corresponding sequence data. While named species make up a small percentage of eukaryotic diversity (**Fig. 1A**), these data provide a valuable window into current biases in research efforts.

**Figure 1.**
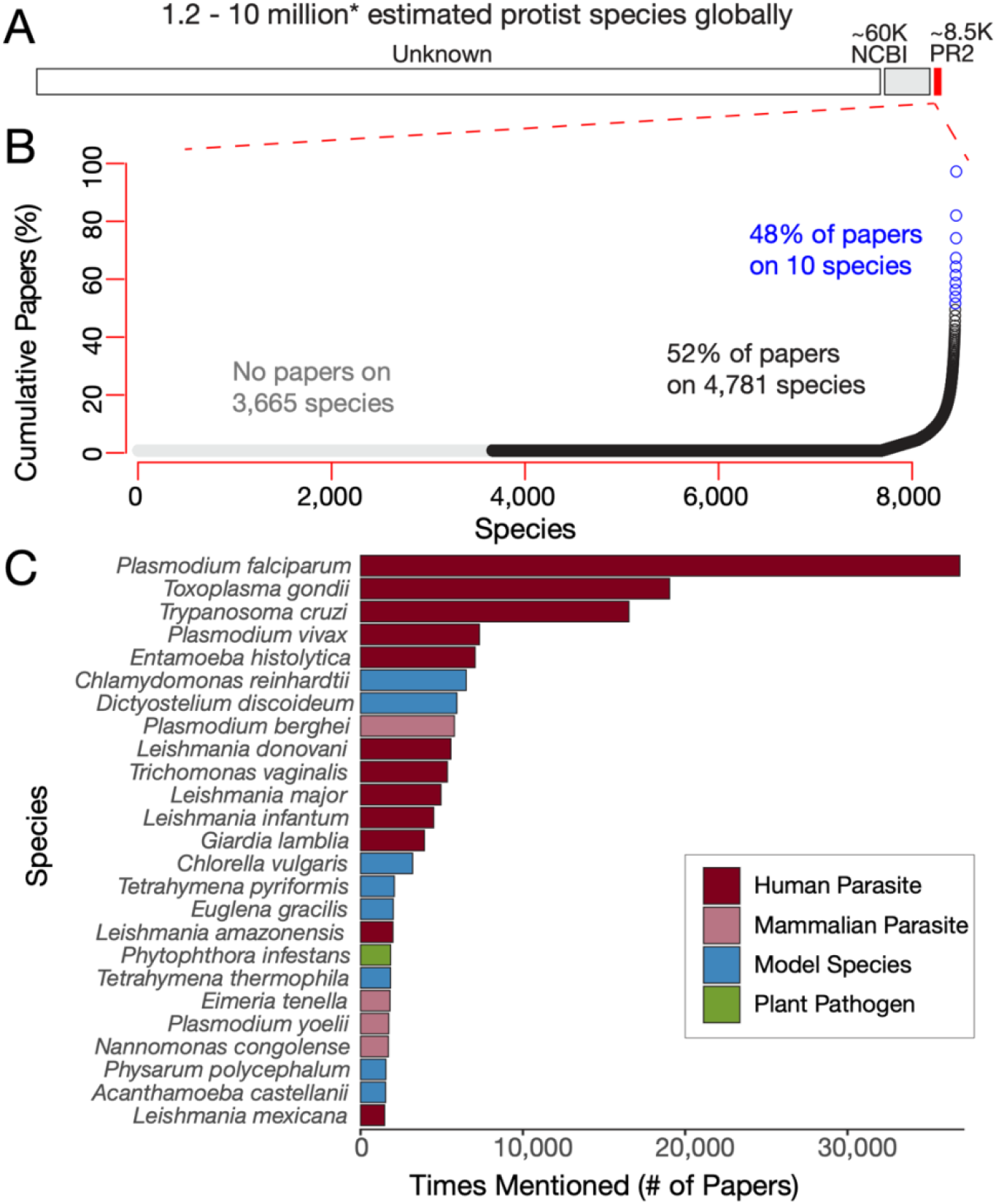
The vast majority of microbial eukaryotic research is focused on a small number of species, mostly human and mammalian parasites. A) There are an estimated 1.2-10 million species of protists. Of that, NCBI acknowledges about 60 thousand^16^. The PR2 database, used in this study, has about 8.5 thousand unique species of protists. B) The top 10 protists make up 48% of the papers in PubMed, and 3,665 species remain unstudied. C) The distribution of papers on protists is skewed, with the top 25 making up 62.67% of all protist publications. Each colored bar represents the number of papers each protist is mentioned in. The color of the bar indicates the function of each species, showing that the most studied type is human parasites.

We found that the top ten species comprise nearly half (48%) of the literature on microbial eukaryotes. Further, the top 25 protist species make up nearly two-thirds of all published research (63%; **Fig. 1B**). The most frequently mentioned species are primarily human parasites. Twelve parasites (including *Plasmodium falciparum, Toxoplasma gondii, Trypanosoma cruzi*, and *Plasmodium vivax*) collectively make up 75% of the top 25 species and 47% of all PubMed literature on protists. *Plasmodium falciparum*, the parasite responsible for malaria in humans, was the top referenced protist, mentioned in 39,915 papers and comprising 15% of the total literature. In contrast, the 15th most-mentioned protist, *Tetrahymena pyriformis*, makes up less than 1%. Notably, 43% of the species in PR2 were not mentioned in a single research article on PubMed. Beyond parasites, the top 25 species also included eight lineages considered nonparasitic model species for protists, such as *Chlamydomonas reinhardtii* and *Dictyostelium discoideum* (**Fig. 1C**). We note that even most microbial eukaryotic model species are far less developed than *Escherichia coli* or *Saccharomyces cerevisiae*, with limited or no genetic tools available^17^. Although both *E. coli* and *C. reinhardtii* (the ‘model’ protist with the most mentions on PubMed) were developed as model organisms in the 1940s^18,19^, the difference in accumulated knowledge between them is drastic. *E. coli* has over 300,000 hits on PubMed while *C. reinhardtii* has only 6,504 hits and lags far behind in genetic tool development and characterization of its basic biology and interactions with other microbes. Research on protists is much less widespread than that of bacteria and other organisms. Indeed, there are more mentions of *E. coli* than the totality of protists. As a result, our understanding of microbial eukaryotes, even some of the most well-studied, lags significantly behind that of other microbial groups.

### Bias in sampling environments

We observed a large discrepancy between the sampling sources of the most frequently sequenced protists versus those most frequently discussed in the literature (**Fig. 2**). Although research efforts on protists are dominated by mammalian parasites found in blood (**Fig. 2A**), the most frequently sequenced protists were from aquatic habitats including both marine and freshwater environments (**Fig. 2B**). Sequenced species were mostly free-living, often planktonic, and included several clades of poorly described dinoflagellates (Dinophyceae, Dino Group I Clade 1, Group II Clade 6, etc.), diatoms (Bacillariophyceae, Mediophyceae), and ciliates (Sessilida, Oligotrichida, Strombidiidae). Our findings demonstrate that the habitats from which protists are most often collected and sequenced does not translate to those most studied in the literature. In other words, the lack of references to these groups is not a result of insufficient sampling.

**Figure 2.**
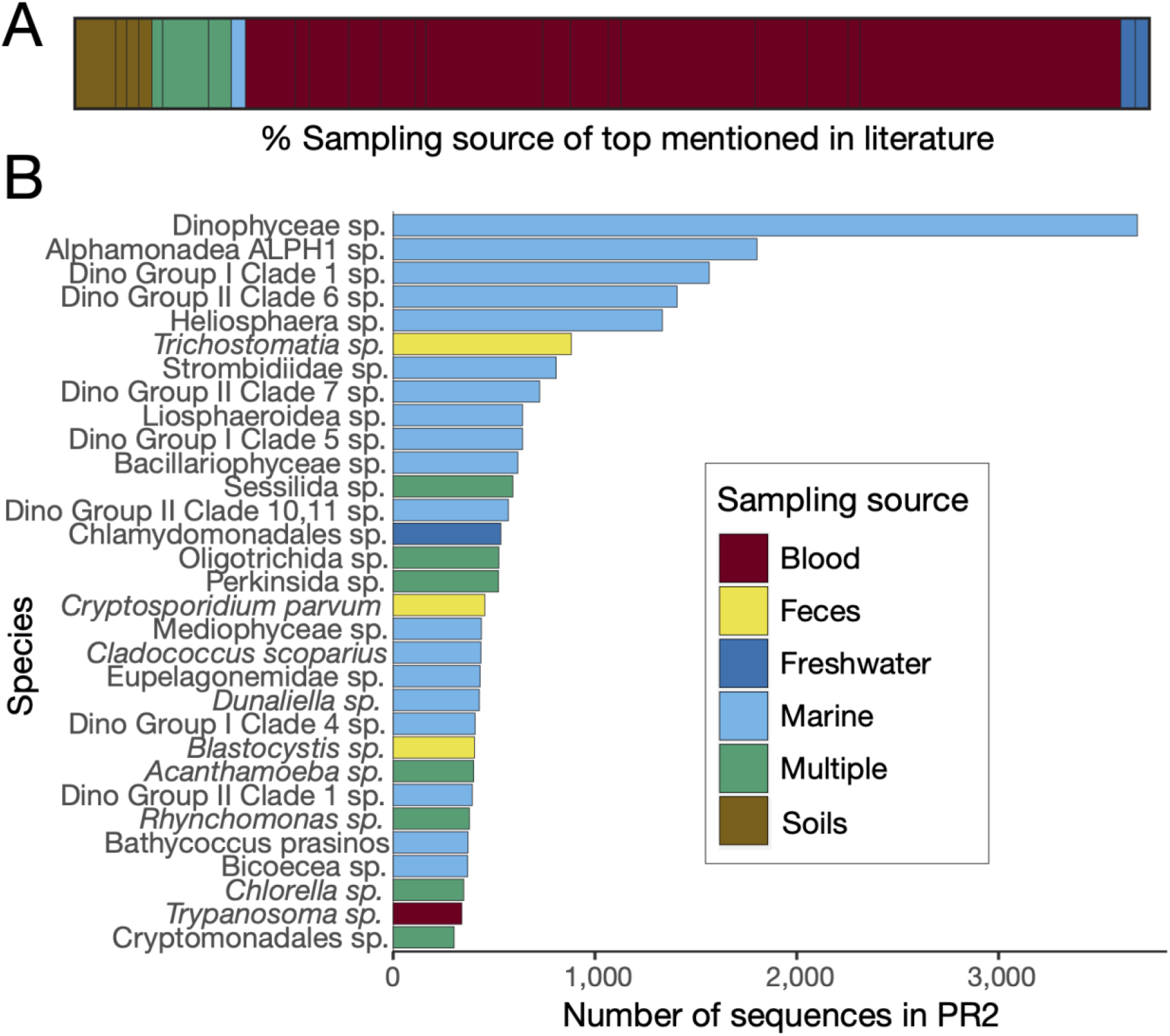
There is a mismatch between the sampling source of protists most mentioned in literature versus the sources where protists are most frequently sampled. A) A bar plot summarizing the top mentioned species on PubMed. Black bars represent distinct species and are scaled by relative percent of mentions. Colors represent the reported sampling source (including blood, feces, freshwater, marine, soils, and multiple sources reported). B) A bar plot displaying the top sequenced species in the PR2 Database. The most frequently sampled species are primarily sourced from marine environments, and nearly all are aquatic, with only a few found in feces or blood. The ‘Multiple’ category is comprised of species sampled from multiple environments.

It is important to note that the most sequenced protists also reflect a bias in sampling of aquatic environments relative to other habitats, namely terrestrial soils, in which protists are known to be abundant and diverse. None of the top 30 protists by sequence abundance were sampled from soil, despite soil environments likely hosting orders of magnitude more species diversity than aquatic environments^2,20^. This underrepresentation is significant, as soil protists play key roles as primary producers, decomposers, and even pathogens^21^ directly contributing to fertility, plant growth, and overall ecosystem function^22,23^. Other habitats such as extreme environments (for example, geothermal springs, glaciers, deep-sea vents, arid deserts) that have long been a focus for bacterial and archaeal sampling also remain poorly sampled^24^.

Microbial compendium and genome initiatives have yielded a wealth of information on the bacteria and archaea that inhabit diverse environments, but characterization of microbial eukaryotic genomes from complex environments lags greatly behind^25^. However, advances in long-read sequencing, enrichment techniques, and bioinformatics pipelines designed for microbial eukaryotes present exciting opportunities to narrow these gaps in ecosystem-level ‘omics characterization^26^ and rapidly advance our understanding of protists from poorly sampled ecosystems.

### Phylogenetic diversity offers opportunities for expanded research

To contextualize research effort of protists in a phylogenetic framework, we built a tree of unique protist sequences in PR2 and mapped the number of corresponding mentions in PubMed across the phylogeny. We found several major clades including Stramenopiles, Dinoflagellates, and Rhizaria with a large number of distinct sequences but which had no representatives in the list of the top 20 most mentioned protists. The stramenopiles exhibit a remarkable ecological and functional diversity, from oxygen-producing algae and kelp to pathogenic oomycetes. Intriguingly, they also represent the group with the highest proportion of unstudied species (**Supplementary Fig. 1**), highlighting a key target for future research. Though it is known that stramenopiles play key roles in oxygen production, a lack of cultured strains inhibits our knowledge of their specific roles and contributions^27^. Another major clade for which we detected a large number of environmental sequences but that remains poorly studied is Rhizaria. For example, recent work in Foraminifera (*Allogromia laticollaris*) has demonstrated that individuals spend a majority of their time in an endoreplicated stage, challenging the paradigm that genomes predominately cycle between haploid and diploid states^28^. It is likely that additional characterization of species within these clades will continue to yield new insights on cellular and genomic biology of eukaryotes.

Even the best studied lineages are within clades of mostly unknown or poorly characterized species. Apicomplexa (within Alveolata) makes up a majority of the top studied species, such as *Plasmodium* species and *Toxoplasma gondii* (**Fig. 3A**). Yet other major clades within Alveolata including dinoflagellates and ciliates have many more sequences than apicomplexans yet are referenced far less (**Fig. 3B**; over 90 thousand times compared to 12 thousand for ciliates). Dinoflagellates may be of interest due to their unique genome structure and gene regulation^29^. Ciliates have already challenged the rules of biology with widespread non-canonical genetic codes and unusual genomic architecture^30,31^. Increased characterization efforts leveraging cutting-edge methods (such as RNA interference (RNAi) tools, high throughput proteomics, and cryo-electron microscopy) can help their progress in rapidly becoming tractable models^32^.

**Figure 3.**
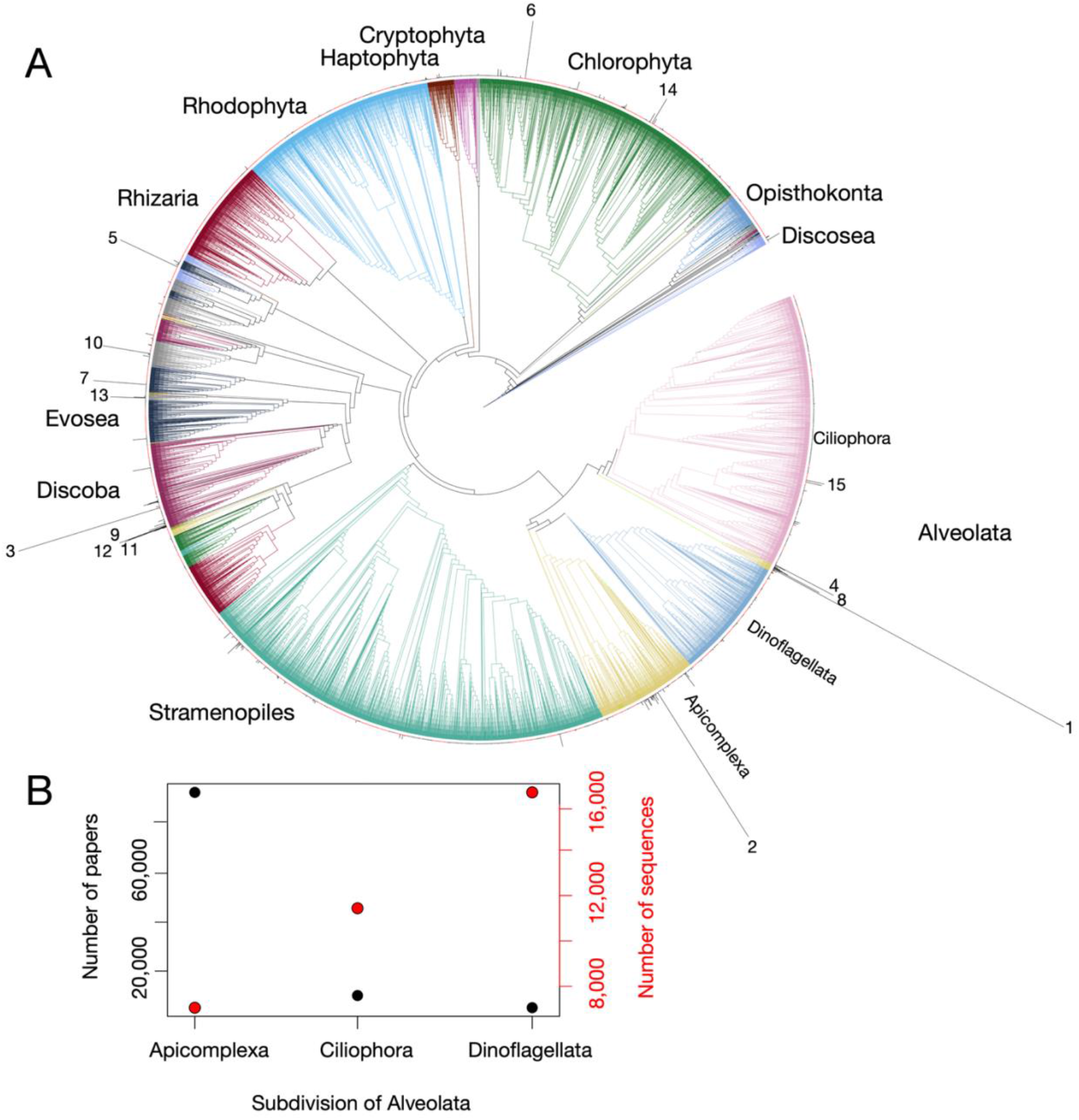
Major clades of microbial eukaryotes have not been the focus of a scientific article, including most parasite diversity. A) Phylogenetic tree of 8,470 18S rRNA sequences from protists in PR2. Sequences were aligned with mafft and the tree was built with RAxML-HPC BlackBox on CIPRES. The tree was visualized on iToL, with branches colored by major clades. Length of black bars extending from each tip is proportional to the number of times each organism is mentioned in PubMed. Some clades are unresolved due to short sequences in PR2. B) Looking into the Alveolata division reveals specific bias. Red dots represent the number of sequences in the PR2 Database for each subdivision, and black dots represent the number of papers the species of each subdivision were mentioned in.

We also quantified the relative research effort of fungal species (one lineage nested within the major clade Ophistokonta) compared to all microbial eukaryotic lineages (**Supplementary Fig. 2A**). Although fungi represent only a small fraction of eukaryotic phylogenetic diversity, these charismatic organisms have long been a research focus and are well-appreciated for the key roles they play in ecosystems. Not surprisingly, *S. cerevisiae* and *Candida albicans* (**Supplementary Fig. 2B**) both have a greater number of mentions than any protist. *S. cerevisiae* has 38,227 more mentions than *P. falciparum*, the most mentioned protist. The total number of mentions (for the 8,118 fungal species) was 320,031, whereas the total for protists was only 242,844. Only 14 protists were in the top 25 species when fungi were added in, underscoring the unbalanced research emphasis.

### Outlook

Recognized for their human health consequences, parasites have overwhelmingly received the most research among protists, but they are a small minority of protist species. Parasites are also poorly representative of eukaryotic diversity, often with reduced genomes and fast evolutionary rates^33,34^. We can aim to reduce research bias to better understand eukaryotic diversity, yet it is unrealistic to expect that every known protist species can be comprehensively examined. This raises an important question: how should we determine which species merit focused research, beyond those with clear relevance to human health? We propose a few guiding criteria to focus on for the in-depth characterization of morphological, physiological, molecular, and genomic diversity of protist lineages. A first obvious target is coverage of major clades that lack a representative species that has been cultured, described morphologically and physiologically, and for which no genome is available. Second, we also suggest another consideration should be species with significant contributions to ecological or environmental health of environmental systems. Finally, we argue that cross-disciplinary research at both the ecosystem-scale and at the molecular and cellular level will be critical to discovery and elucidation of novel protists that challenge the rules of eukaryotic life. Description of unknown lineages across understudied ecosystems is a key source of discovery for lineages that do biology differently while development of genetic tools in emerging model systems are critical to the characterization of novel biology.

## Methods

To compile a list of protists from the PR2 Database, plants, animals, and fungi were filtered out, as well as any samples that did not have an 18S rRNA sequence. Separately, fungi were added back in to compare the research effort of fungi relative to protists. To first look at bias in literature, the number of papers for each species were found by using the PubMedR package in Rstudio. Full names (e.g. *Plasmodium falciparum*) were searched for in Titles and Abstracts to find the number of papers associated with a species, as some species were common phrases (e.g. ammonia) and some abbreviated names (e.g. *A. acuta*) were duplicated. Analyses were completed in R via ggplot and base R packages. Lists of the unique species with each corresponding number of papers for protists and fungi are on figshare. In order to highlight sampling bias, locations of samples from the PR2 Database were referenced. The location descriptions were not normalized, and not every sample had a listed source. A list of locations provided by the PR2 database is on figshare. To look at phylogenetic bias, a single representative entry was chosen for each species and species name, division, and 18S rRNA sequences were exported from the PR2 database. This resulted in 8,470 sequences (and one outgroup sequence), which were run through mafft alignment with default parameters, trimmed via the bioconda package trimAl^35^, and then used to build a phylogenetic tree via RaxML-HPC BlackBox algorithm on CIPRES Science Gateway^36,37^, and visualized on iTOL (Interactive Tree of Life^38^. Figures were finalized using Adobe Illustrator to alter color, spacing, and text labels. A cartoon tree (modified from ref. 39), was modeled to offer context in the diversity of protists and their evolutionary proximity to fungi.

## Funding

J.A.L. acknowledges support from the Syracuse University Office of Undergraduate Research & Creative Engagement (SOURCE Bridge Award). H.B.R. was supported by the NSF Graduate Research Fellowship. A.M.O. was supported by grants from the U.S. National Science Foundation (DEB #2439029), the National Aeronautics and Space Administration (Exobiology #80NSSC25K7916), and the American Philosophical Society Lewis and Clark Fund for Exploration and Field Research in Astrobiology.

## Data availability

All code and data used in the analyses is available at: https://doi.org/10.6084/m9.figshare.c.7862267

## Supplementary Information

**Supplementary Figure 1.**
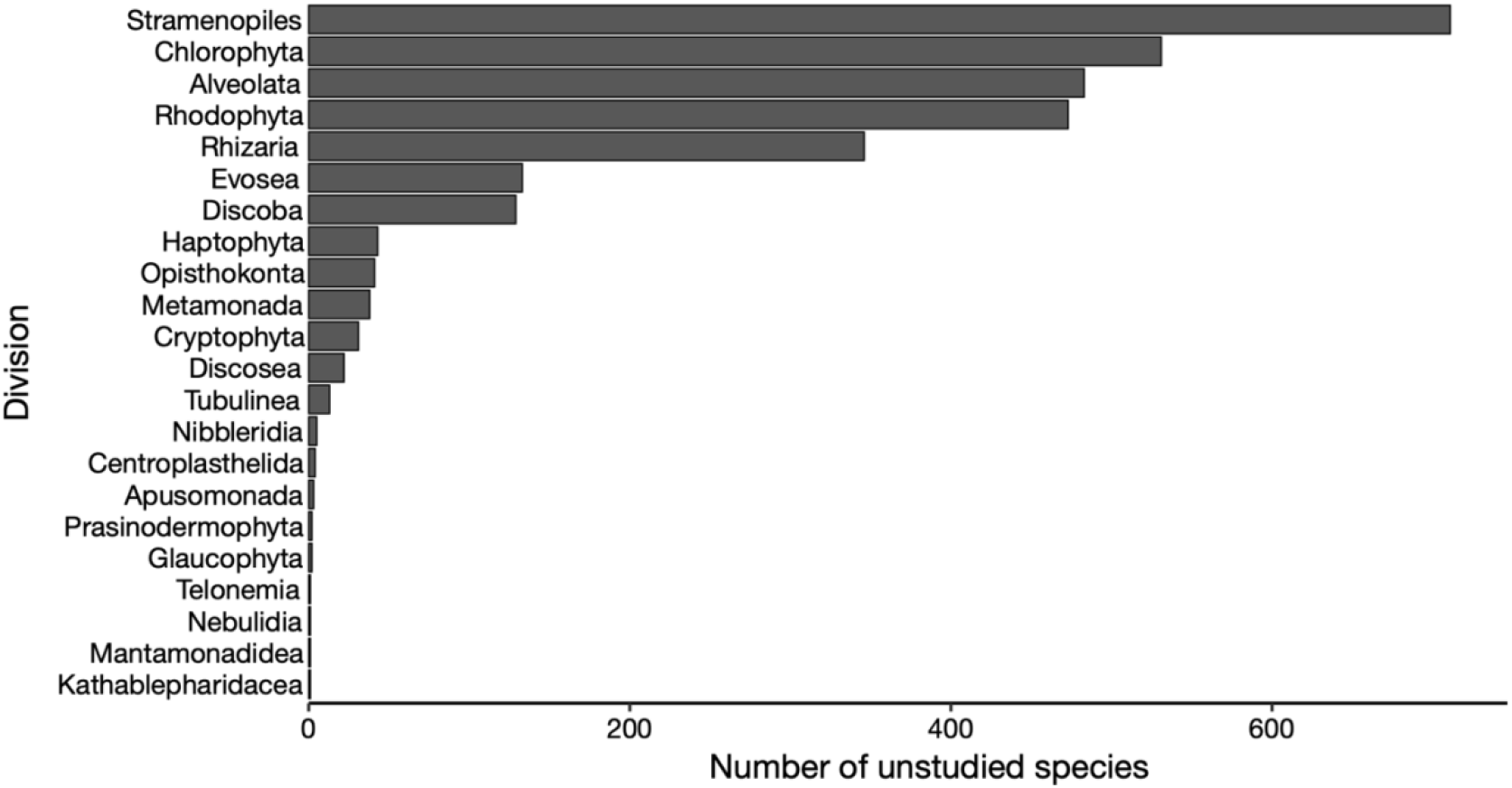
Clades with the greatest number of named species that were not mentioned in any PubMed articles. Bars represent the number of unique species in each clade from the PR2 database that were found in zero articles on PubMed.

**Supplementary Figure 2.**
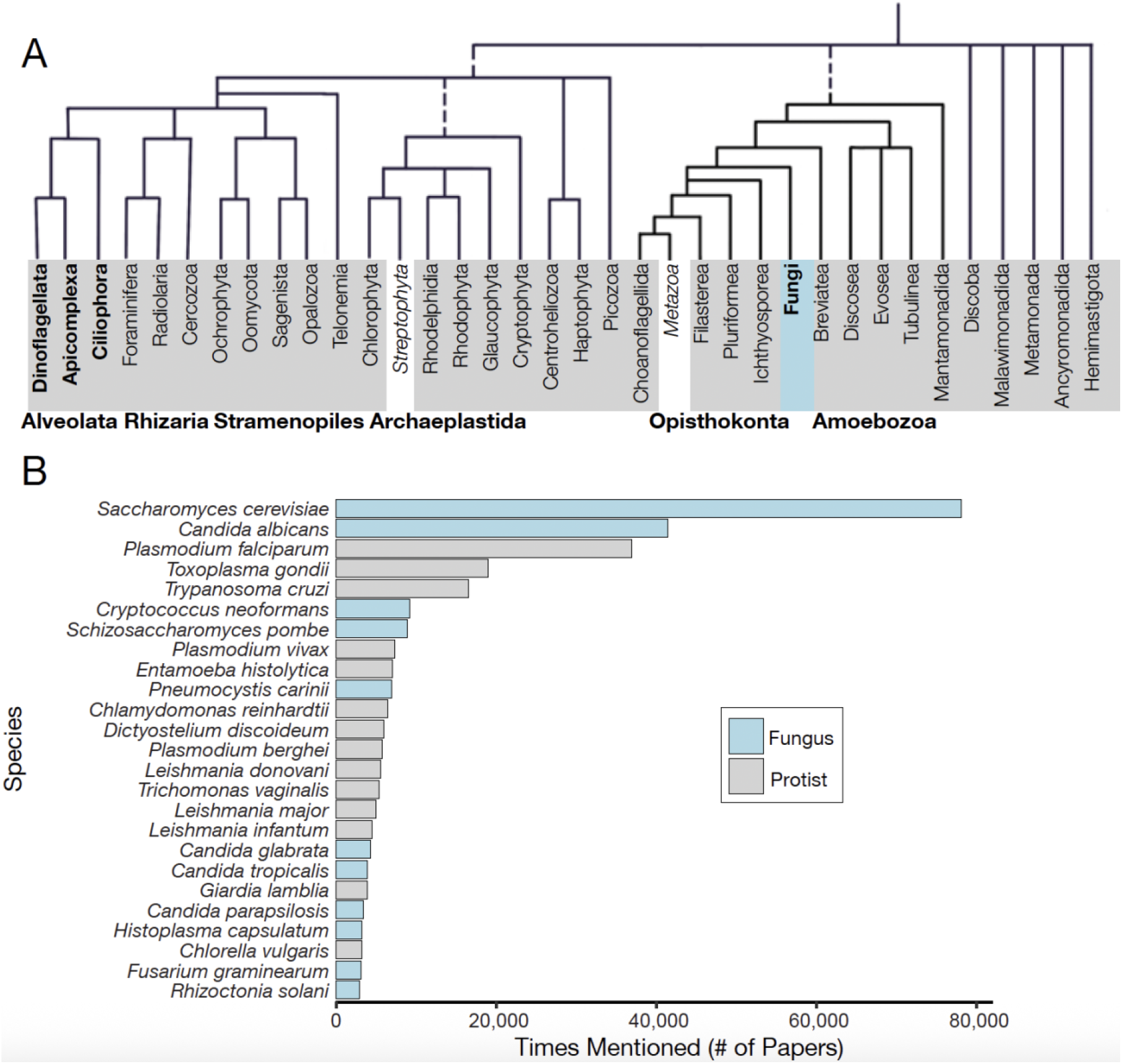
Fungi displace the top protist, but protists keep the majority of the top 25 places. A) Cartoon tree of Eukaryotes. Fungi is highlighted in the same blue as Part B to act as a reference for the difference in diversity. Modified from ref. 24, adapted from refs. 39–40. B) Bar plots reflecting the top 25 microbial eukaryotes mentioned in literature when fungi are included in counts. Plants and animals are still excluded from this assessment. The PR2 database was also used for searching fungal species.

**Table S1.** The vast majority of microbial eukaryotic research is focused on human and mammalian parasites. Table of the top 15 protists, sorted in descending order by number of articles mentioned on PubMed.

| Species | Number of Papers | Percentage | Category |
| --- | --- | --- | --- |
| *Plasmodium falciparum* | 39,915 | 15.20 | Human parasite |
| *Toxoplasma gondii* | 19,037 | 7.84 | Human parasite |
| *Trypanosoma cruzi* | 16,531 | 6.81 | Human parasite |
| *Plasmodium vivax* | 7,322 | 3.02 | Human parasite |
| *Entamoeba histolytica* | 7,058 | 2.91 | Human parasite |
| *Chlamydomonas reinhardtii* | 6,504 | 2.68 | Model species |
| *Dictyostelium discoideum* | 5,929 | 2.44 | Model species |
| *Plasmodium berghei* | 5,778 | 2.38 | Mammalian parasite |
| *Leishmania donovani* | 5,562 | 2.29 | Human parasite |
| *Trichomonas vaginalis* | 5,347 | 2.20 | Human parasite |
| *Leishmania major* | 4,951 | 2.04 | Human parasite |
| *Leishmania infantum* | 4,504 | 1.85 | Human parasite |
| *Giardia lamblia* | 3,932 | 1.62 | Human parasite |
| *Chlorella vulgaris* | 3,205 | 1.32 | Model species |
| *Tetrahymena pyriformis* | 2,076 | 0.85 | Model species |

## Notes

https://doi.org/10.6084/m9.figshare.c.7862267

### Competing Interest Statement

The authors have declared no competing interest.

